# Disrupted mitochondrial response to nutrients is a presymptomatic event in the cortex of the APP^SAA^ knock-in mouse model of Alzheimer’s disease

**DOI:** 10.1101/2024.02.02.578668

**Authors:** Andrés Norambuena, Vijay Kumar Sagar, Zhuoying Wang, Prakash Raut, Ziang Feng, Horst Wallrabe, Evelyn Pardo, Taylor Kim, Shagufta Rehman Alam, Song Hu, Ammasi Periasamy, George S. Bloom

**Author notes:** Corresponding to: Andrés Norambuena, Department of Biology, University of Virginia, PO Box 400328, Charlottesville, VA 22904-4328, USA.

## Abstract

**Introduction:** Reduced brain energy metabolism, mTOR dysregulation, and extracellular amyloid-β oligomer (xcAβO) buildup characterize AD; how they collectively promote neurodegeneration is poorly understood. We previously reported that xcAβOs inhibit <u>N</u>utrient-induced <u>M</u>itochondrial <u>A</u>ctivity (NiMA) in cultured neurons. We now report NiMA disruption *in vivo*.

**Methods:** Brain energy metabolism and oxygen consumption were recorded in APP^SAA/+^ mice using two-photon fluorescence lifetime imaging and multiparametric photoacoustic microscopy.

**Results:** NiMA is inhibited in APP^SAA/+^ mice before other defects are detected in these amyloid-β-producing animals that do not overexpress APP or contain foreign DNA inserts into genomic DNA. GSK3β signals through mTORC1 to regulate NiMA independently of mitochondrial biogenesis. Inhibition of GSK3β with lithium or TWS119 stimulates NiMA in cultured human neurons, and mitochondrial activity and oxygen consumption in APP^SAA^ mice.

**Conclusion:** NiMA disruption *in vivo* occurs before histopathological changes and cognitive decline in APP^SAA^ mice, and may represent an early stage in human AD.

## Introduction

It is well accepted that reduced brain energy metabolism is a key feature of AD. Clinical studies tracking 2-deoxy-2-[^18^F] fluoro-d-glucose (FDG)-PET uptake, a surrogate for glucose utilization in brain, support the hypothesis that brain hypometabolism is one of the earliest AD biomarkers and is detectable years before symptom onset [1]. Although FDG-PET screening has provided valuable information allowing identification of patients at risk of AD and other brain disorders, the molecular mechanism(s) at fault remain poorly understood.

The amyloid cascade hypothesis [2] has guided much of our understanding about mechanisms of AD pathogenesis and most drug discovery efforts for AD [3–5]. Key players in the hypothesis are Aβ peptides, ∼40 amino acid long proteolytic fragments of the amyloid precursor protein (APP) and the principal amyloid plaque component. Although plaques, along with neurofibrillary tangles composed mainly of tau are the defining histopathological lesions in AD, a preponderance of evidence indicates that the Aβ and tau species most responsible for the synapse loss and neuron death that together cause AD symptoms are soluble oligomers that accumulate in brain pre-symptomatically [6].

A few years ago, we reported a novel form of communication between lysosomes and mitochondria, a fundamental cell biological process that we named NiMA as an acronym for <u>N</u>utrient-induced <u>M</u>itochondrial <u>A</u>ctivity [7]. More specifically, NiMA represents activation of the master regulatory protein kinase, mTORC1 [8], by insulin, or arginine and leucine (R+L), to trigger mitochondrial respiration within minutes in the absence of mitochondrial biogenesis. Importantly, we also found that xcAβOs block NiMA in cultured neurons [7]. The mechanism involves xcAβO-mediated ectopic activation of mTORC1 at the neuronal plasma membrane [9] and upregulation of superoxide dismutase 1 (SOD1), a major regulator of cellular redox and a pathogenic factor in ALS [10]. Whether NiMA is disrupted in live AD brain, however, was not explored.

GSK3β has been studied for several years in the context of AD [11]. It is a major tau kinase [12], and studies in mouse models of AD strongly implicate hyperactive GSK3β in AD pathogenesis [13]. Additionally, GSK3β has been linked to the control of mitochondrial anterograde trafficking [14], and feeding mice lithium, which inhibits GSK3β [15,16], was shown to modulate brain energy metabolism [17].

Using two-photon fluorescence lifetime imaging (2P-FLIM) and multi-parametric photoacoustic microscopy (MP-PAM), we tracked mitochondrial activity and brain energy metabolism in the cortex of wild type (WT) and heterozygous APP knock-in mouse littermates. The heterozygous knock-ins, henceforth called APP^SAA/+^, express one copy of the endogenous WT mouse *APP* gene, with the second endogenous *APP* gene modified by homologous recombination to include a humanized Aβ_1-42_ region containing the Swedish (KM670/671NL), Arctic (E693G), and Austrian (T714I) mutations. In contrast to nearly all other APP transgenic mouse strains, APP^SAA^ mice do not overexpress APP, nor do they include any foreign transgene inserts that might toxically interrupt one or more endogenous mouse genes. The pathological progression of APP^SAA^ mice has been extensively characterized [18], allowing correlation of brain energy metabolic changes to disease progression.

We now show that NiMA inhibition occurs long before the accumulation of plaques, increased CSF tau, microgliosis, and cognitive decline recently reported for APP^SAA/+^mice [18]. Mechanistically, we found that GSK3β controls mitochondrial activity in neurons in culture and live mouse brain by a mechanism involving a GSK3β-mTORC1 signaling pathway. Importantly, pharmacological inhibition of GSK3β also stimulated mitochondrial activity and oxygen consumption in the cortex of the APP^SAA/+^ mice. Thus, disruption of NiMA in brain could be a very early event in AD pathogenesis in humans.

## Materials and Methods

### Cell Culture and materials

Human embryonic kidney cells (Lenti-X 293T Cell line from Takara. Cat no. 632180) were grown in DMEM/F12 media (GIBCO) supplemented with HyClone cosmic calf serum (GE Healthcare) and 50 µg/ml gentamycin (GIBCO).

Human neurons were differentiated from ReNcell VM neuronal precursor cells (EMD Millipore) as described earlier [7]. For 2D differentiation, the cells were plated into 35 mm glass-bottom dishes with DMEM/F12 differentiation media (GIBCO) supplemented with 2 μg/ml heparin (ScienceCell), 2% (v/v) B27 neural supplement (GIBCO) and 50 µg/ml gentamycin (GIBCO) without growth factors. One-half volume of the differentiation media was changed every 3 days until the cultures were used for experiments [19].

### Mouse Strain

The hAbeta^SAA^ (APP^SAA^ KI) knock-in mouse model of Alzheimer’s disease was purchased from The Jackson Laboratory (Strain #:034711). Animals were housed in a pathogen-free barrier facility with a 12-hour light/12-hour dark cycle, and *ad libitum* access to food and water under a protocol approved by the IACUC of the University of Virginia. All pups were genotyped by q-PCR following The Jackson Laboratory’s recommendations. All experiments, except for assays shown in Supplemental Figure 2, were performed in males heterozygous for APP^SAA^ KI or WT littermates.

### Lentivirus production and infection

Lentiviral particles for shGSK3β knockdowns were prepared as follows. The expression plasmids, pLKO.1 (Mission shRNA library from Sigma-Aldrich, see below), and the packaging vectors, pSPAX2 and pMD2.G (Addgene plasmids 12260 and 12259, respectively) were transfected using Lipofectamine 3000 (ThermoFisher) into HEK293T cells grown in 15 cm Petri dishes to ∼80% confluence in DMEM (GIBCO) supplemented with 10% HyClone cosmic calf serum. Each transfection was with 15 µg total DNA at a 50%/37.5%/12.5% ratio of expression vector/pSPAX2/pMD2G. Lentivirus-conditioned medium was collected 24 and 48 hours after the start of transfection. Lentiviral particles were concentrated in a Beckman Coulter Optima XE-90 ultracentrifuge for 2 hours at 23,000 rpm (90,353 g_max_), at 4° C in an SW32Ti rotor, resuspended in 400 µl Neurobasal medium and stored at −80° C in 20 µl aliquots. Cultured neurons were transduced and incubated for 72 hours before assays were performed.

### Human shRNA Sequences

The following plasmids were purchased from The RNAi Consortium of the Broad Institute; 1) TRCN0000039564 (#1) targeting sequence CATGAAAGTTAGCAGAGACAA and TRCN0000039565 (#2) targeting sequence AGCAAATCAGAGAAATGAAC. These constructs are already cloned into pLKO.1, and lentivirus particles were prepared as described in a previous section.

### Mouse Brain Cell Extracts

Brain cortex samples were placed in a Wheaton 1ml Dounce homogenizer containing 1ml of lysis buffer to cover the sample. Lysis buffer consisted of RIPA buffer (Bioworld, #42020024-2); 1% HALT™ protease inhibitor cocktail; 1% phosphatase inhibitor cocktail 2 (Sigma Aldrich, #P5726); and 1% phosphatase inhibitor cocktail 3 (Sigma Aldrich, #P0044). Brain homogenates were placed in an Eppendorf 5415C centrifuge at 10,000 rpm (8,160 g_max_) for 10 minutes at 4° C. Finally, supernatants were transferred to new tubes and used as protein lysates for western blots.

### Antibodies

The following antibodies were from Cell Signaling Technologies: rabbit anti-GSK3β, catalog number 9832 (dilution 1:3,000); mouse anti-GSK3β phosphorylated on Serine 9, catalog number 9323 (dilution 1:3,000); mouse anti-S6 ribosomal protein, catalog number 2317 (dilution 1:3,000); and rabbit anti-S6 ribosomal protein phosphorylated on Serine 240/244, catalog number 5364. Antibodies from Proteintech included: mouse anti-PGC1α, catalog number 66369-1-1g (dilution 1:3,000); and mouse anti-TOM40, catalog number 66658-1-1g (dilution1:5,000). Mouse anti-human APP (clone 6E10) was from Biolegend (cat no. 803014; dilution1:5,000). Mouse anti-total OXPHOS rodent antibody cocktail was from Abcam (catalog number ab110413; dilution 1:10,000).

### Immunoblotting

Samples were resolved by SDSPAGE using either 10 or 12% acrylamide/bis-acrylamide gels and transferred to 0.22 µm nitrocellulose (Bio-Rad). Membranes were blocked with Odyssey blocking buffer (LI-COR Biosciences) and were incubated with primary antibodies and secondary IRDye-labeled antibodies diluted into antibody buffer (Odyssey blocking buffer diluted 1:1 with PBS/0.1% Tween 20). All antibody incubations were overnight at 4° C, and 3 washes of 5 minutes each with PBS/0.1% Tween 20 were performed after each antibody step. Membranes were quantitatively scanned by a Odyssey imaging station (LI-COR Biosciences).

### Multiparametric Photoacoustic Microscopy (MP-PAM)

All animal procedures were approved by the Institutional Animal Care and Use Committee at Washington University in St. Louis. Following hair removal, a surgical incision was made on the scalp. The exposed skull was cleaned. Then, the mouse skull region over a 3×3 mm^2^ ROI was carefully removed to expose the cortex for topical application of Leucine (0.8mM) plus Arginine (0.4mM) or TWS119 (100μM; Cayman Chemicals, catalog no. 10011251). After craniotomy, the anesthetized mouse was transferred to the MP-PAM system. The animal body temperature was maintained at 37 °C via a heating pad and the local brain temperature was also maintained at 37°C via a temperature-controlled water tank. Ultrasound gel was applied between the open-skull window and water tank for acoustic coupling. Following the baseline imaging, the ultrasound gel was gently removed and a solution of either Leucine (0.8mM) plus Arginine (0.4mM) or TWS119 (100μM) was applied topically. The exposed mouse cortex was treated with the corresponding solution for up to 80 minutes. Then, the ROI was covered again with ultrasound gel and subjected to post-treatment MP-PAM imaging [20]. MP-PAM uses dual-wavelength (i.e., 532 nm, 558 nm) and high-repetition-rate nanosecond-pulsed laser excitation to achieve simultaneous pixel-wise measurement of the oxygen saturation of hemoglobin (sO_2_) and cerebral blood flow speed. In the acquired MP-PAM images, vessel segmentation was performed and the average values of sO_2_ and flow speed in main arteries and veins within the FOV are calculated in the vessel segments. Oxygen extraction fraction (OEF) is measured by calculating the relative difference in arterial and venous sO_2_.

### NADH and NADPH measurements

### In vivo 2P-FLIM imaging

#### Time-correlated single photon counting (TCSPC) Fluorescence Lifetime Imaging Microscopy (FLIM)

Surgical preparation of animals and application of amino acids or TWS 119 were done as described in the previous paragraph. Live 2P-FLIM was recorded on a Zeiss LSM-780 NLO confocal/multiphoton microscopy system comprising an inverted Axio Observer Z1 microscope, an X-Cite 120PC Q mercury arc light source (Excelitas Technologies) for cell selection, a motorized stage for automated scanning, an IR Chameleon Vision-II ultrafast Ti:sapphire laser for multiphoton excitation (Coherent), a Zeiss 40X 1.3 NA oil immersion Planapo objective, an environmental chamber (PeCon GmbH, Germany) that envelops the microscope stage to control temperature and CO_2_ level, and a 3-channel FLIM system based on three HPM-100-40 GaAsP-based hybrid detectors and 3 SPC-150 TCSPC boards (Becker & Hickl). The SPC-150 boards are synchronized with the 2-photon excitation laser and the Zeiss LSM-780 NLO scan head signal. Ex 740nm; Em450/50 nm.

#### Imaging

Human neuronal cultures were grown in 35-mm glass-bottom dishes, maintained at 37° C in 5% CO_2_/95% air on the stage of the Zeiss LSM-980 NLO microscope. Cultures were developed in 20-25 days. Then, cells were treated with the indicated inhibitor for ∼80 minutes at 37° C. The laser was tuned to 740 nm with an average power of 7 mW at the specimen plane, and NAD(P)H fluorescence was collected using a 450-500 nm emission filter (Objective lens, 40x). For each experiment, 10 fields of view were recorded in the descanned mode, and then each field of view was subjected to a 40-second acquisition in the non-descanned mode. The laser power and acquisition time were selected to ensure enough photons per pixel while avoiding photodamage to cells.

#### Processing

FLIM images were processed with SPCImage software (v5; Becker & Hickl) at maximum-likelihood fitting. All FLIM image parameter data were exported for further processing in Fiji software (https://hpc.nih.gov/apps/Fiji.html). Here, thresholded, normalized photon images selected pixel-ROIs from the more prominent intensity of the mitochondrial morphology by a standard Fiji plugin. Mitochondrial ROIs were applied to all FLIM parameter data creating a mitochondria-specific data pool by a custom Fiji plugin. Those results were analyzed by a custom code in Python software, where all parameters were first examined and filtered for obvious outliers (insignificant numbers). The FLIM metric of interest – a_2_%, the fraction of enzyme-bound NAD(P)H – an indicator for changes in the mitochondrial metabolic OXPHOS state was extracted. NAD(P)H-a_2_% data was plotted for publication-ready charts and statistics in Microsoft Excel.

### Statistics

The paired t-test was used for analyzing all 2P-FLIM assays. Since thousands of ROIs were obtained per image, the average ROI for each field of view was calculated to reduce the sample size and thus the number of false positives. All data were assumed to be normally distributed. For MP-PAM assays, vessel segmentations and quantitative analysis were done following our established protocol [20]. The paired t-test was used to compare the cerebral hemodynamics and oxygen metabolism before and after the application of experimental compounds. A *p*-value of less than 0.05 was considered statistically significant.

## Results

### Transient decrease in brain metabolism in APP^SAA^ mice

We first measured baseline mitochondrial activity in the cortex of 2, 4 and 6-month-old APP^SAA^ mice, and by comparison, in WT littermates. To do this, we used 2P-FLIM to monitor NADH and NADPH fluorescence lifetimes, and the degree to which these co-enzymes associate with partner enzymes in mitochondria [7]. While the enzyme-bound fraction of NADH, expressed as “a_2_%” [21], is a surrogate for mitochondrial respiration [7], the corresponding a_2_% measurement for NADPH reports activation of biosynthetic pathways [22]. Although the excitation and emission spectra of NADH and NADPH are identical, 2P-FLIM allows measuring their distinct fluorescence lifetimes [22]. Accordingly, we determined the relative contributions of each coenzyme to the total measured fluorescence lifetimes under baseline conditions. This approach revealed that on average, NADH accounted for 55% of the observed fluorescence lifetime signals, while the remaining 45% was due to NADPH. An ∼19% reduction in NADH and a corresponding increase in NADPH were observed in 2 month old APP^SAA/+^ mice when compared to WT littermates, but these differences did not persist as animals aged up to 6 months old (Figure 1A-C).

**Figure 1.**
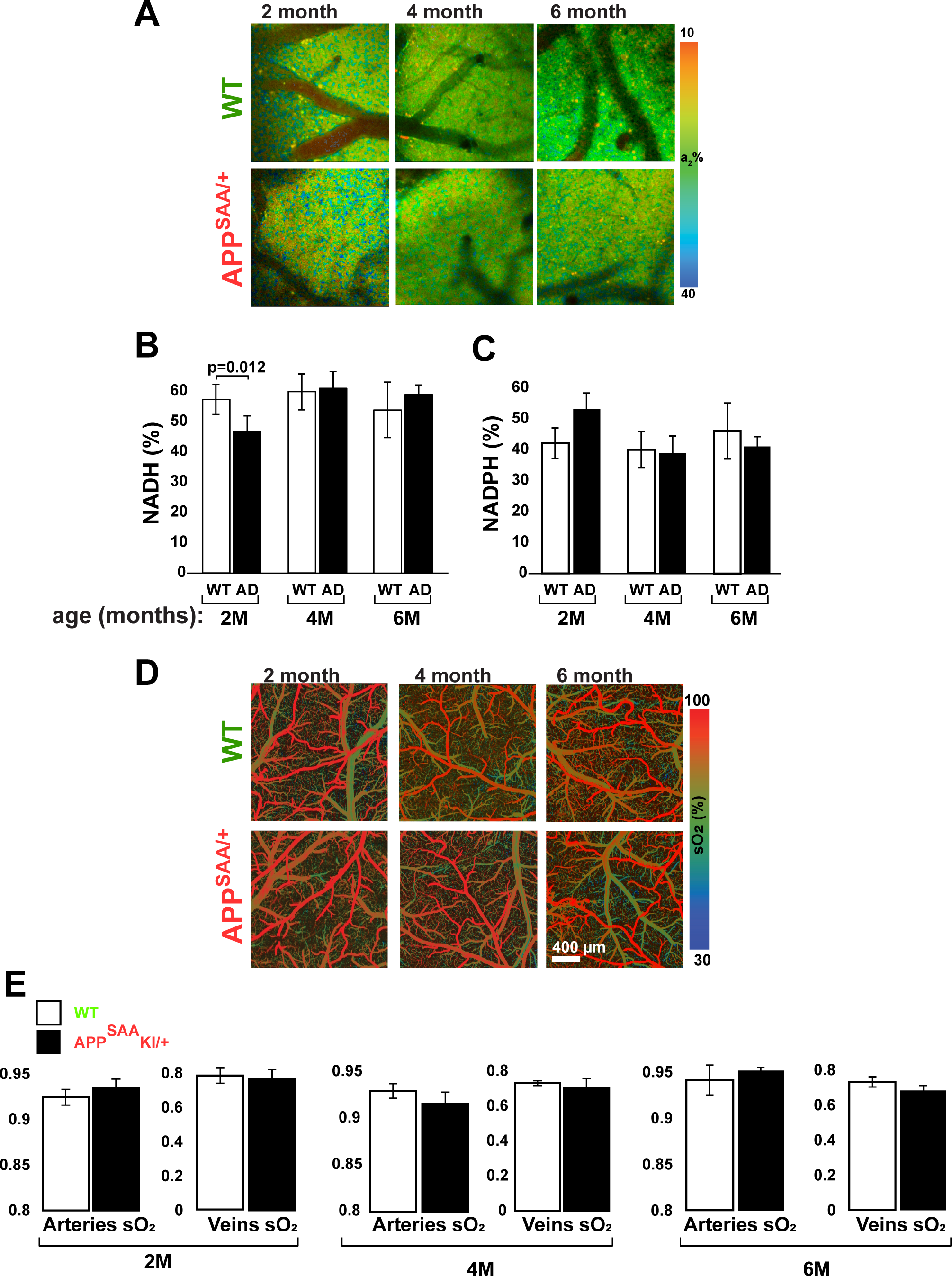
2P-FLIM and MP-PAM intravital imaging shows similar cortical energy metabolic activity in WT and APP^SAA/+^ mice. A-C, 2P-FLIM imaging at baseline revealed a transient ∼19% decrease in enzyme-bound NADH and the corresponding increase in enzyme-bound NADPH in 2-month-old APP^SAA/+^ mice. D-E, MP-PAM imaging at baseline revealed virtually identical levels of mitochondrial respiration irrespective of age or genotype. 4-6 animals per group were used for NADH/NADPH measurements, and 4 animals per group were used for MP-PAM. Student’s two-tailed unpaired t-tests were used to assess statistical significance, and error bars represent +/- s.e.m.

In parallel experiments, we applied MP-PAM to evaluate baseline levels of oxygen consumption in live mouse brain [20]. MP-PAM imaging through a cranial window enables simultaneous, high-resolution detection of total hemoglobin concentration (CHb), oxygen saturation of hemoglobin (sO_2_), and cerebral blood flow. Notwithstanding the transient decrease in NADH detected by 2P-FLIM, sO_2_ levels in the resting cortex were virtually constant irrespective of age or genotype (Fig 1D-E).

We also analyzed the expression levels of key proteins of the electron transport chain (ETC) in 2, 4 and 6 month old mice. Western blots of cortical samples from WT and APP^SAA/+^ animals revealed no age-dependent changes in the expression levels of NDUFB8, SDHB, UQCRC2, MTCO1, or ATP5A, all of which are subunits of corresponding complexes I to V of the ETC (Supplemental Figure 1). Because APP^SAA/+^ mice do not develop amyloid plaques until ∼16 months of age, but do have low levels of Aβ in brain, CSF, and plasma by 4 months [18], these results suggest that a low concentration of Aβ causes a transient decrease in energy metabolism that does not compromise steady-state oxidative phosphorylation (OXPHOS) in the cortex. The data are consistent with observations made more than 30 years ago in human brain that mitochondrial activity is normal at early stages of AD [23]. APP^SAA/+^ mice can thus serve as a credible model for exploring mechanisms that regulate brain energy metabolism and might go awry in AD.

### Early disruption of mitochondrial response to amino acids in APP^SAA/+^ mouse cortex

We recently reported NiMA inhibition in cultured mouse and human neurons exposed to xcAβOs, and in human neurons that were not exposed to xcAβOs but were co-cultured with otherwise identical human neurons that express the K670N/M671L (Swedish) and V717I (London) APP mutations, and secrete Aβ [7,10]. To seek evidence for NiMA inhibition *in vivo*, we used 2P-FLIM in the current study to compare mitochondrial activity in cortex of WT versus APP^SAA/+^ mice. We topically applied R+L to the cortex through an open skull window, which caused the expected increase in OXPHOS (a_2_%) in 2 month old WT mice. Increased mitochondrial activity was detected by 30 minutes after application of the amino acids, persisted for up to 80 minutes, and occurred regardless of the animals’ age (Figure 2A-C). In contrast, when the same approach was used for APP^SAA/+^ mice, the mitochondrial response to R+L was observed at 2 months (Figure 2D) but was completely absent in 4 (Fig 2E), 6 (Fig. 2F) and 8 month old (not shown) animals. In addition, the mitochondrial responses to amino acids in 4 month old mice were not influenced by gender (Supplemental Figure 2).

**Figure 2.**
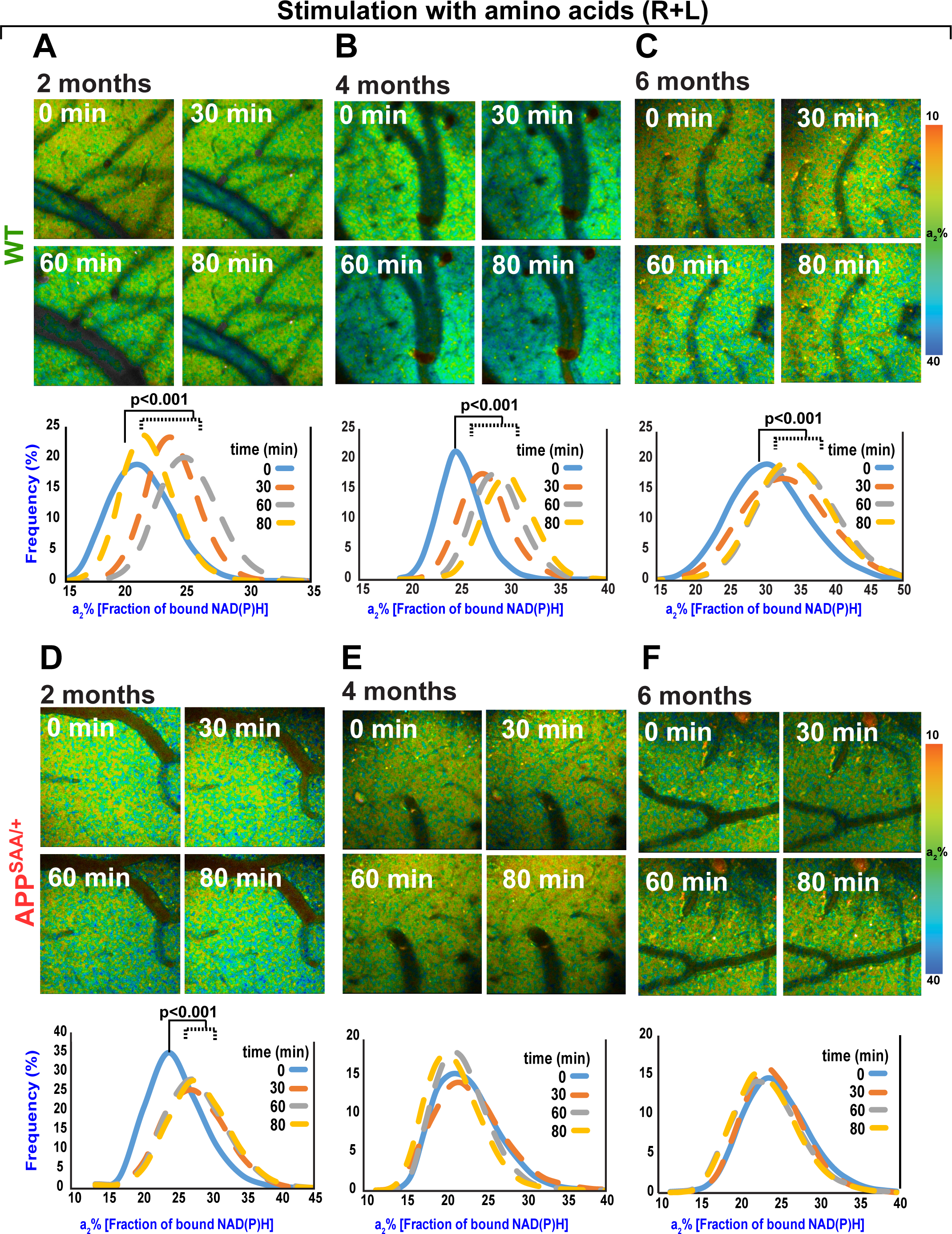
2P-FLIM imaging reveals disruption of mitochondrial response to amino acids in APP^SAA/+^ mouse brain. A-C, 2P-FLIM imaging of WT mouse cerebral cortex through an open-skull window after topical application of amino acids (R+L). An increase in a_2_%, indicating upregulation of mitochondrial activity, was observed 30 minutes after the initial application of amino acids and remained elevated for up to 80 minutes. A similar response was observed regardless of the age of the animals tested. D-F, 2P-FLIM imaging of APP^SAA/+^ mice showed an increase in a_2_% in response to amino acids only in the 2 month old group. Statistical analyses were performed using Student’s two-tailed unpaired t-test. 3-4 animals/age/genotype were used in these experiments.

In parallel experiments, we used MP-PAM to test if fluctuations in a_2_% reflect changes in mitochondrial respiration in mouse brain. Topical application of R+L through an open skull window reduced sO_2_ in WT mice by ∼10 and 15% in arteries and veins, respectively, (Figure 3), suggesting an increase in oxygen consumption due to mitochondrial activity in tissue surrounding blood vessels. These changes were similar in magnitude irrespective of the age of the animals (Figure 3). When MP-PAM microscopy was applied to APP^SAA/+^ mice, we observed that amino acids triggered a comparable reduction in sO_2_ in 2 month old animals, but not in older mice (Figure 4). Altogether, this *in vivo*, label-free imaging of mitochondrial energy metabolism in APP^SAA/+^ mice suggests that low levels of soluble Aβ peptides inhibit NiMA in the cortex.

**Figure 3.**
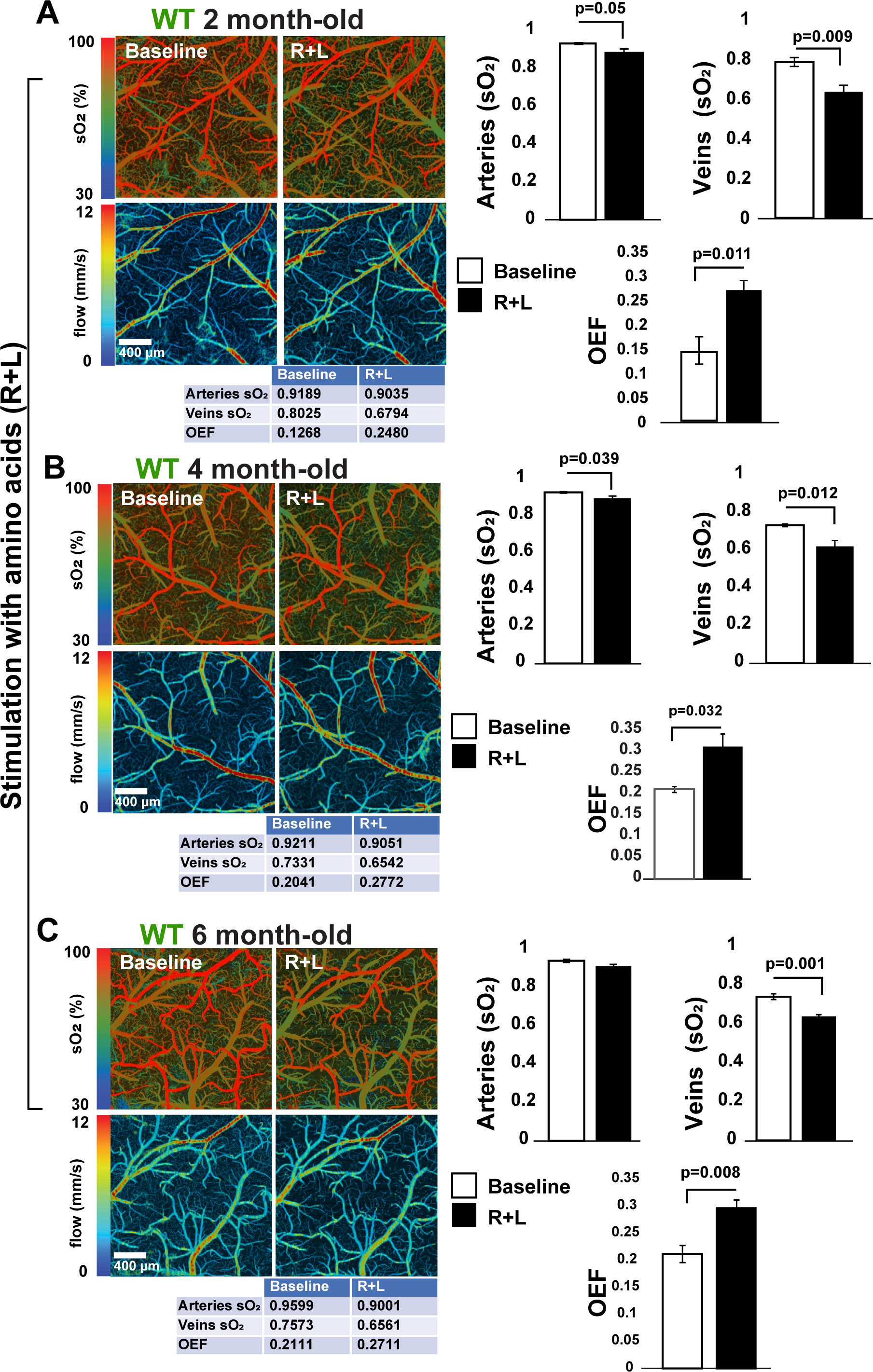
Amino acids (R+L) trigger oxygen consumption in WT mouse brain. A-C, MP-PAM imaging of WT mouse cerebral cortex through an open-skull window before and 80 minutes after topical application of arginine and leucine (R+L). A decrease in sO_2_ in the cortical vasculature was observed at all ages, indicating elevated oxygen extraction and consumption due to the upregulation of mitochondrial activity. sO_2_ in arteries, veins, and Oxygen extraction fraction (OEF) values for the experiments shown in the figure are depicted in the table below each image. Bar graphs show quantification of 4 independent experiments. Statistical analyses were performed using Student’s two-tailed unpaired t-test. Error bars represent +/- s.e.m. 4 animals/age group were used.

**Figure 4.**
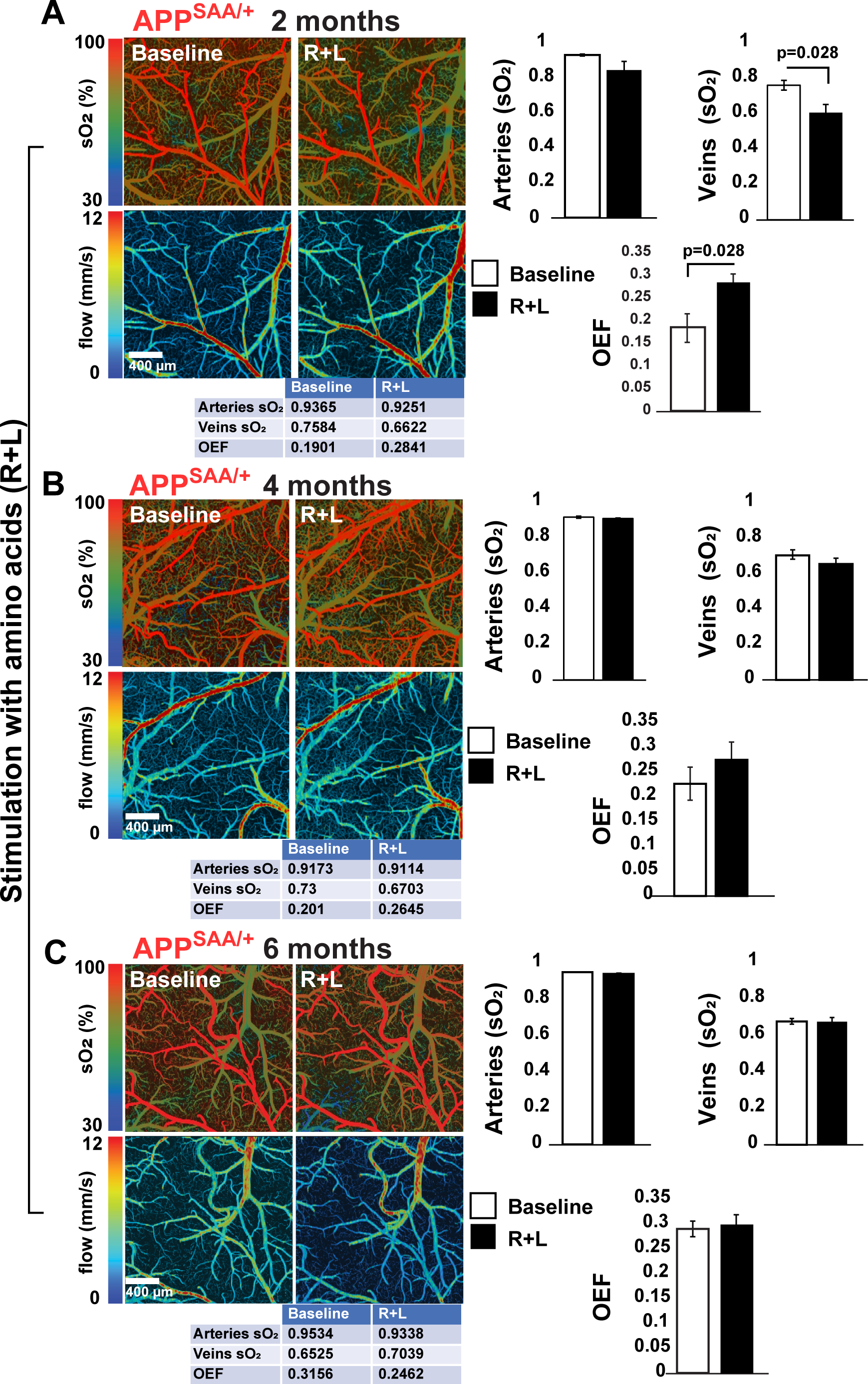
Amino acid-induced oxygen consumption is blocked in APP^SAA/+^ mouse brain. A-C, MP-PAM imaging of APP^SAA/+^ mouse cerebral cortex through an open-skull window before and 80 minutes after a topical application of amino acids (R+L). Decreases in cortical vasculature sO2 and oxygen extraction fraction (OEF) were observed only in the 2 month old group. sO_2_ in arteries, veins, and Oxygen extraction fraction (OEF) values for the experiments shown in the figure are depicted in the table below each image. Bar graphs show the quantification of 4 independent experiments. Statistical analyses were performed using Student’s two-tailed unpaired t-test. Error bars represent +/- s.e.m. 4 animals/age group were used in these experiments.

### GSK3**β** regulates NiMA in neurons in culture and live mouse brain

To test if GSK3β controls mitochondrial functioning, human neurons differentiated from the ReNcell VM line of human neuronal progenitors [7] were first treated for 80 minutes with increasing concentrations of TWS119, a GSK3β inhibitor [24]. Mitochondrial activity was monitored by 2P-FLIM of neurons exposed to the inhibitor or a vehicle control. The contributions of enzyme-bound NADH and NADPH to the total fluorescence lifetime was extracted and plotted as described earlier [7,22]. TWS119 treatment at 500nM had no effect until 18 hours (not shown), but 1 and 10μM TWS119 increased the proportion of NADH over NADPH in a concentration-dependent manner within 80 minutes (Figure 5A), indicating an increase in mitochondrial respiration. Under these experimental conditions, we observed the expected increase in phosphorylation of serine-9 on GSK3β (Figure 5B), which inhibits its kinase activity [25]. Inhibition of GSK3β did not change the level of PGC1α, a major regulator of mitochondrial biogenesis (Figure 5B) and was paralleled by increased phosphorylation of the ribosomal S6 protein, a downstream effector of mTORC1 and a read-out mTORC1 activity (Figure 5B), suggesting that GSK3β controls NiMA through activation of mTORC1. We also tested if Li+, another well-known GSK3β inhibitor [15,16] triggered similar effects. As shown in Supplemental Figure 3, treating human neuron cultures with LiCl increased both the fraction of NADH and the phosphorylation of GSK3β on Ser-9. Then, we verified whether TWS119 increases mitochondrial activity in the live mouse brain. 2P-FLIM assays revealed that topical application of TWS119 for 80 min through a cranial window also caused a ≥30% increase in a_2_% in WT mouse brain (Figure 5C).

**Figure 5.**
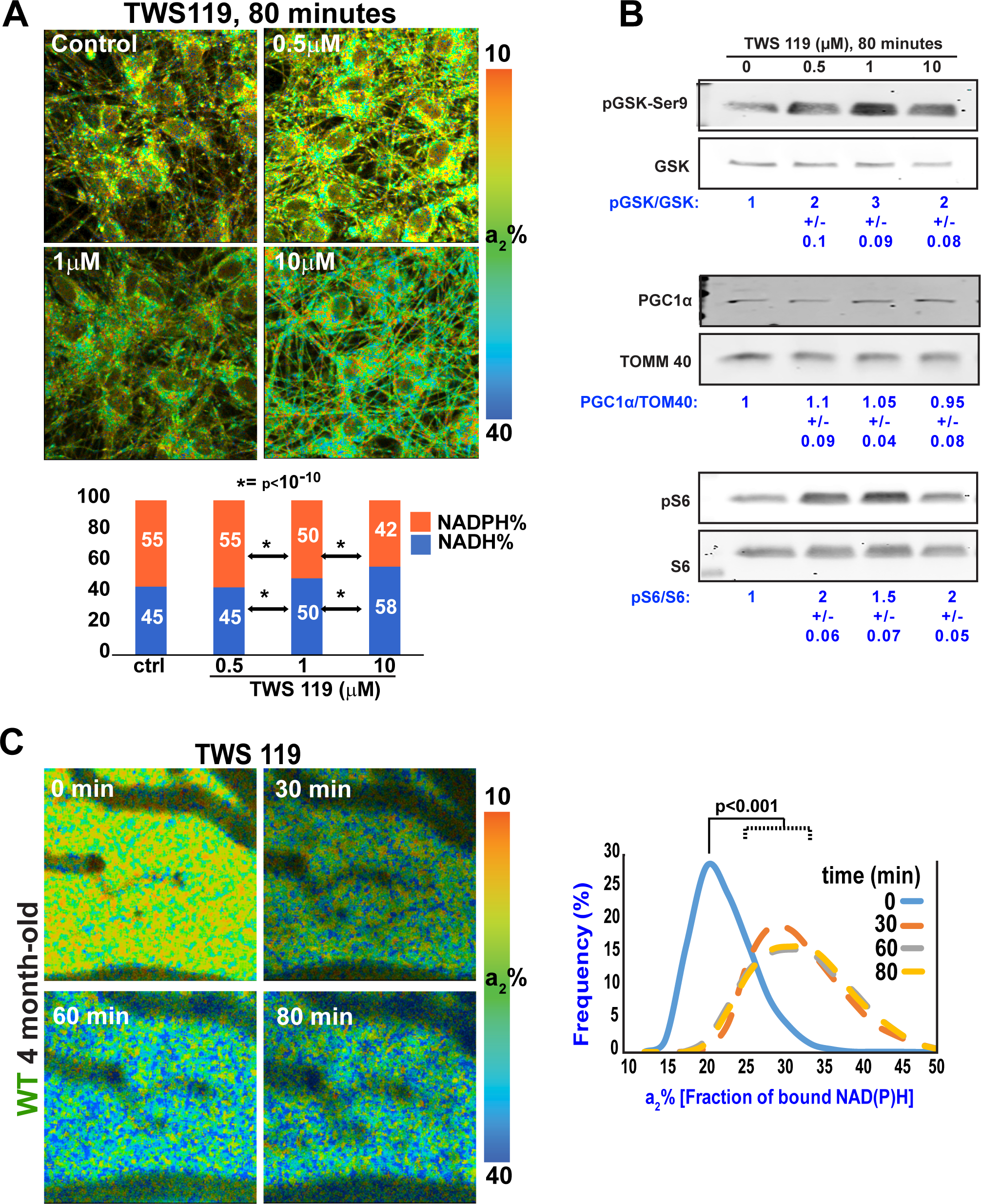
GSK3β inhibition stimulates mitochondrial activity in cultured human neurons and WT mouse brain. A, Neurons derived from the RenCell VM line of human neuronal progenitors were differentiated for 20 days. Then, one culture was left untreated (control) while others were treated with TWS119 at the indicated concentration for ∼80 minutes. 2P-FLIM revealed that inhibition of GSK3β increased the fraction of bound NADH, indicating a rise in mitochondrial respiration. Micrographs are representative of 3 independent experiments. B, TWS119 efficiency was tested by western blots of protein samples from the same experiments used for 2P-FLIM. TWS119 caused large increases in phosphorylation of the inhibitory site, serine-9 on GSK3β, and in phosphorylation of the ribosomal protein S6, a readout of mTORC1 activity. Changes in mitochondrial activity occurred without changes in expression of the mitochondrial transcription factor PGC1α. The experiment is representative of 3 independent assays. C, 2P-FLIM imaging of WT mouse cerebral cortex through an open-skull window after topical application of TWS119. An increase in a2% was observed for several minutes indicating upregulation of mitochondrial activity. Statistical analyses were performed using Student’s two-tailed paired t-test. 4 animals were used in these experiments.

To corroborate that GSK3β was mediating the effect of TWS119 and lithium on mitochondrial respiration, we used lentiviral-driven delivery of antisense shRNA to knock down GSK3β by ∼50% in human neurons. As we found for TWS119 (Figure 5) and LiCl (Supplementary Figure 3), GSK3β knockdown increased the NADH/NADPH ratio in cultured human neurons (Figure 6A-C). Similar to TWS119 (Figure 5B) and LiCl (Supplemental Figure 3B), GSK3β knockdown did not change the level of PGC1α (Figure 6D), fortifying our previously reported evidence that NiMA activation does not require an increase in mitochondrial number [7]. Collectively, the results for TWS119, LiCl and GSK3β knockdown establish that GSK3β regulates mTORC1 activity in neurons, and extend our prior finding that stimulation of lysosomal mTORC1 increases NADH and OXPHOS in neuronal mitochondria [7]. GSK3β is thus a newly identified upstream regulator of NiMA in neurons.

**Figure 6.**
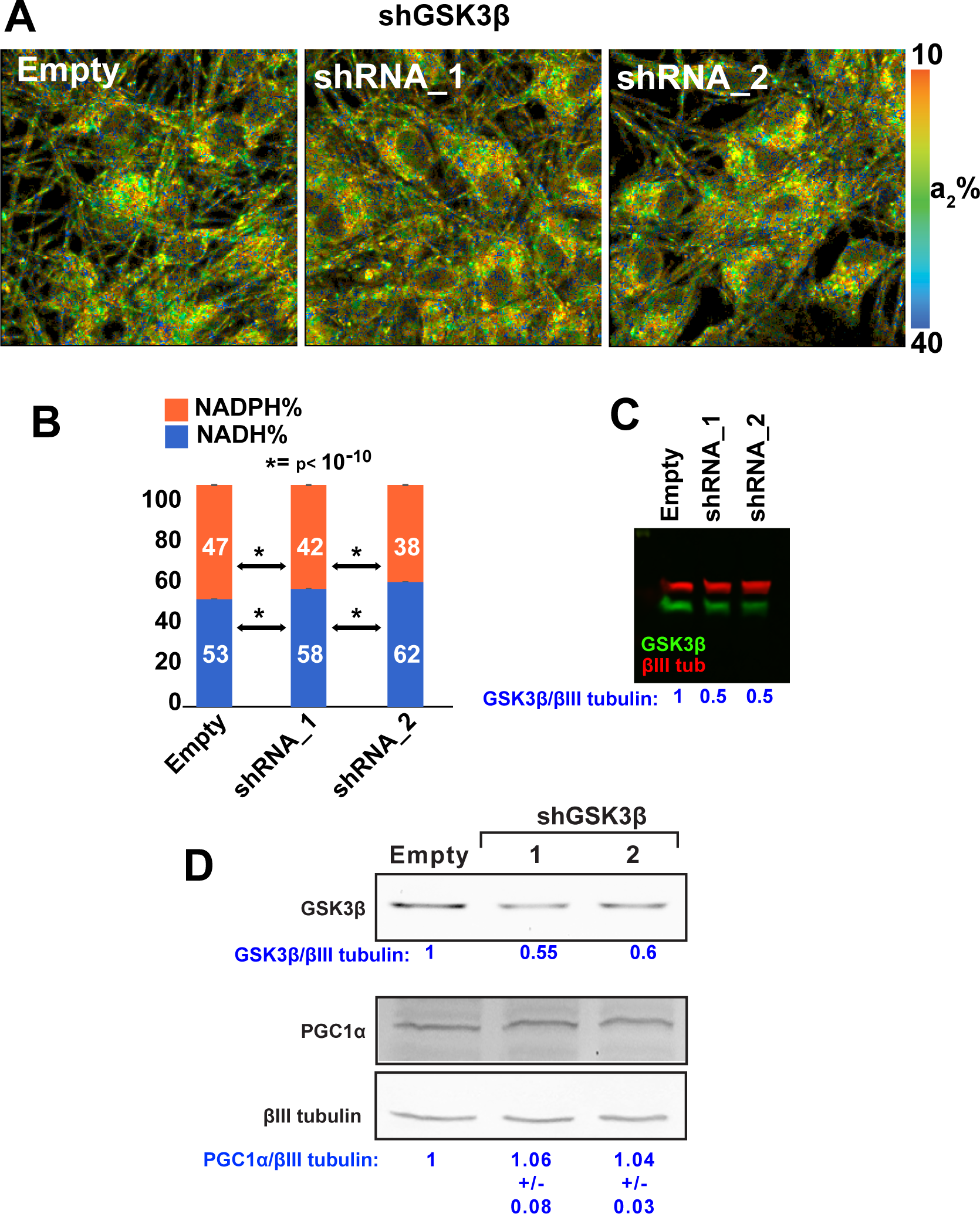
GSK3β reduction stimulates mitochondrial activity in cultured human neurons. A-B, GSK3β expression was reduced by lentiviral-mediated delivery of antisense shRNA. 2P-FLIM analysis in live cells revealed an increase of NADH, indicating a rise in mitochondrial respiration in GSK3β knock-down cells. The experiment is representative of 3 independent assays. C-D, GSK3β knockdown efficiency in the 2P-FLIM assays was ∼50%. Under similar experimental condition, there were no changes in the expression of PGC1α. The experiment is representative of 3 independent assays.

### TWS119 rescues mitochondrial activity in the APP^SAA^ KI/+ mice

We and others have shown that AβOs impair insulin/IGF1 signaling in neurons and insulin resistance is a major risk factor for developing AD [26]. Mechanistically these effects are partially explained by a mechanism including (i) AβOs-induced internalization of IR receptors [27], (ii) reduction in AKT activity, a downstream IR target [10], and (iii) increased association with lysosomes of the tuberous sclerosis complex, and mTORC1 inhibitor [9]. As AβOs also affect the anterograde trafficking of mitochondria through activation of GSK3β in a tau-dependent manner [14], it is possible to speculate that AβOs-induced and tau-dependent insulin resistance led to GSK3β activation in AD. Thus, inhibiting GSK3β *in vivo* at presymptomatic stages of AD could represent a potential treatment.

As TWS119 increased mitochondrial activity in WT live mouse brain (Figure 5C), we then tested TWS 119’s ability to stimulate mitochondrial activity in the 4-month-old APP^SAA^ KI/+ live brain (when NiMA is unresponsive). Like our observations in WT animals and cultured neurons (Figure 5), TWS119 application for 80 min through a cranial window also increased a_2_% in the APP^SAA^ KI/+ mice (Figure 7A). This change in a_2_% was also paralleled by a partial increase in oxygen consumption as measured by MP-PAM (Figure 7B).

**Figure 7.**
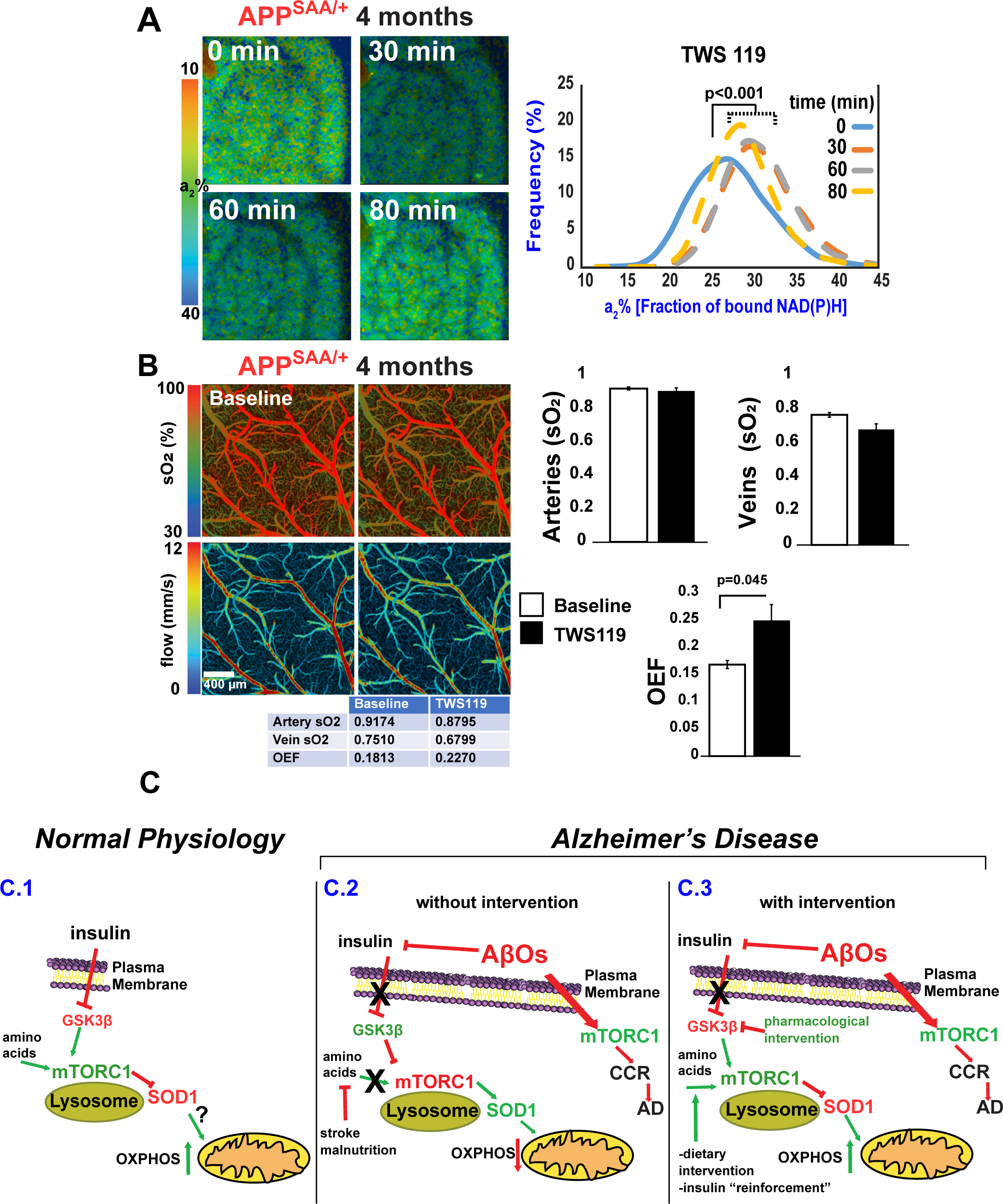
TWS119 triggers mitochondrial respiration in the AD mouse cortex. A-2P-FLIM imaging of the APP^SAA/+^ mouse cerebral cortex through an open-skull window before and after topical application of TWS119. An increase in a2% was observed for several minutes indicating upregulation of mitochondrial activity. Statistical analyses were performed using Student’s two-tailed paired t-test. 3 animals were used in these experiments. B-MP-PAM imaging of APP^SAA/+^ mouse cerebral cortex through an open-skull window before and 80 minutes after a topical application of TWS119. A small decrease in blood oxygenation of the cortical arteries and veins was observed but did not reach statistical significance. Still, there was a significant increase in the overall oxygen extraction fraction. Bar graphs show the quantification of 3 independent experiments. Statistical analyses were performed using Student’s two-tailed unpaired t-test. Error bars represent +/- s.e.m. C-Model for NiMA regulation by GSK3β *in vivo* in brain. C.1, In normal physiological conditions insulin and amino acids stimulate lysosomal mTORC1[7] by a mechanism involving inhibition of GSK3β. Then, mTORC1-mediated phosphorylation of SOD1 at T40 [10], triggers mitochondrial respiration. C.2, Insulin-mediated inhibition of GSK3β is blocked by AβOs, decreasing mTORC1 activity on lysosomes and stimulating it instead at the plasma membrane, where mTORC1 kinase activity triggers cell cycle re-entry (CCR), a frequent prelude to cortical neuron death in AD. C.3, Pharmacological treatments that block AβO-induced activation of GSK3β might restore normal lysosomal mTORC1 activity in AD.

## Discussion

Brain hypometabolism is one of the earliest signs of AD [28–31], and here we present new mechanistic evidence for how this pernicious decline in brain glucose consumption occurs. Using mitochondrial metabolic imaging of live mouse brain, we demonstrate disruption of the NiMA pathway in APP^SAA/+^ mice. The relevance of this process is two-fold. First, this Aβ-driven phenomenon coincides with the earliest detection of soluble Aβ_1-40/42_ in brain and CSF and occurs months before plaques and other key AD features emerge in this transgenic mouse model [18]. Second, NiMA disruption does not impact overall mitochondrial metabolism, which is regulated by a plethora of inputs including a diverse source of nutrients [32], but specifically interferes with mitochondrial respiration triggered by amino acids and likely, insulin as well.

The physiological significance of NiMA inhibition for human AD is underscored by the genotype of the APP^SAA/+^ mice used for this study. APP is expressed in these animals exclusively by the two endogenous murine *APP* genes, one of which is unmodified and the other of which was altered by homologous recombination to contain a humanized Aβ_1-42_ region encoding the Swedish (KM670/671NL), Arctic (E693G), and Austrian (T714I) mutations in place of the corresponding region of WT mouse *APP* [18]. Consequently, and in sharp contrast to nearly all other transgenic APP mouse models of AD, APP^SAA/+^ mice do not overexpress APP, nor do they contain numerous mutant human APP cDNAs that incorporate into mouse genomic DNA at random sites and thereby potentially interfere with normal expression of multiple mouse genes. Just as importantly, the NiMA response to amino acid stimulation is normal in 2 month old APP^SAA/+^ mice, but is completely inhibited by the time animals reach 4 months of age (Figure 2). The simplest explanation for this observation is that the single mutated *APP* gene drove Aβ production to levels sufficient to block NiMA a year before the first plaques appear in these mice at 16 months of age [18]. It follows naturally that NiMA inhibition, which includes reduced ATP production by mitochondria in response to nutrient stimulation, might occur in the brains of human AD patients far in advance of the histopathological features and cognitive decline that define the disease. This soluble Aβ-induced deficiency in the fundamental cellular fuel, ATP, could underly the myriad aspects of neuronal function that deteriorate as AD progresses from its silent, prodromal stage to full blown symptoms.

How does NiMA disruption set the stage for AD pathogenesis? Central to this story are three well-known AD players; AβOs, mTORC1, and GSK3β. Moreover, we previously found that AβO-mediated inhibition of NiMA required the expression of tau [7,10], the building block of neurofibrillary tangles. The physiological mechanisms controlling mTORC1 functions have been widely studied and include central roles for insulin and the amino acids arginine and leucine in activating mTORC1 activity on lysosomes. There, mTORC1 regulates key processes, such as mRNA translation, autophagy, gene expression, and mitochondrial activity [7,8]. At the cellular level, mTORC1-mediated regulation of protein synthesis is required for synaptic function [33] and removal of misfolded proteins and organelles via autophagy [34]. Mitophagy, the removal of defective mitochondria by autophagic mechanisms, is partially regulated by mTORC1 [35] and has been shown to reduce Aβ and tau pathology, and cognitive decline in AD [36]. However, much less is known about the mechanisms leading to mTORC1 dysregulation in AD. Work by our group has shown that AβOs may work as a two-edge sword. At one level, AβOs trigger ectopic activation of mTORC1 at the plasma membrane by a mechanism that is dependent on tau expression, and thereby causes re-entry of neurons into the cell cycle, a prelude to most neuron death in AD [9]. At a second, parallel level, AβOs reduces mTORC1 activity on lysosomes by inhibiting insulin signaling [10]. It is reasonable to speculate that during AD progression the combination of Aβ production and insulin resistance leads to an imbalance in mTORC1 activity favoring its ectopic and toxic activity at the plasma membrane over its normal physiological functions on lysosomes. By extension, these events also lead to NiMA disruption, as we previously showed for cultured neurons [7,10], and as reported in the current study, *in vivo* as well. This hypothesis is consistent with the facts that mTORC1 activity is elevated in AD brain [37], and that decreasing mTOR expression in an AD mouse model ameliorates AD-like pathology and behavior [38]. Thus, NiMA is a fundamental biological process that integrates several key players in AD pathogenesis, including tau, mTOR and autophagy.

The collective findings reported here have therapeutic implications for AD. For example, we show that low doses of lithium, an FDA-approved treatment for mood disorders, increase mitochondrial activity in neurons in culture (Supplemental Figure 3). We also found that TWS119, a GSK3β inhibitor, also increased mitochondrial respiration in both cultured neurons and mouse brain (Figures 5). Just as importantly, we found that TWS119 also increased mitochondrial activity in APP^SAA/+^ mouse cortex (Figure 7A and B). Tideglusib, another GSK3β inhibitor [39] has been tested in clinical trials for AD, and although found to be safe it showed no beneficial effects [40,41]. Interestingly, in our hands, Tideglusib was also ineffective at stimulating mitochondrial respiration in cultured neurons and mouse brain (not shown). Thus, although GSK3β is a promising target for AD therapy, a deeper understanding of its diverse mechanisms of action and cellular effects of GSK3β inhibitors will improve the outcome of future clinical trials directed at this protein kinase.

Several studies have investigated the association between Aβ deposition and local brain energy metabolism, but multimodal FDG-PET and amyloid PET aimed at comparing plaque deposition with changes in local glucose metabolism studies have provided conflicting results [42–48]. Although informative, amyloid PET is limited by resolution and its inability to detect AβOs. Still, amyloid PET studies have suggested that brain areas loaded with plaques are “protected” from Aβ insult [49–51] and represent *in vivo* support for the idea that soluble forms of Aβ are disruptive of cellular energy metabolism. A recent study provided evidence that AβOs are likely initiators of brain hypometabolism [52] and are in line with our findings shown here that AβOs disrupt NiMA *in vivo* in brain.

Many cellular functions disrupted by AβOs are known to be dependent on the expression of tau, [53]. Among them, disruption of synaptic functioning and neurodegeneration have been recognized for a long time, leading to the concept that AD is a “synaptic disease” [54]. The fact that disruption of NiMA was detected in the cortex of APP^SAA/+^ mice months before cognitive decline [18] leads us to propose that chronic metabolic disruption caused by an AβO-induced, tau-dependent mechanism represents a very early event in human AD [55].

## Footnote

Andres Norambuena dedicates this paper to the memory of Edward P. Owens. Ed, during his years of association with the Boom Lab has been most supportive of my work in Alzheimer’s research. I fondly remember his friendly and hopeful outlook that one day we will find a complete answer to this dreadful disease.

## Acknowledgments

We are grateful for financial support from NIH/NIA (grant R01AG067048 to AN); the Owens Family Foundation (GSB); Cure Alzheimer’s Fund (GSB); Rick Sharp Alzheimer’s Foundation (GSB); and the NIH/Office of the Director for funds to purchase a Zeiss 780 and 980 microscopes used in this study (OD016446 and NIH-OD 025156 to AP).

## Conflicts of Interest

The authors have no conflict of interest to report.

## Author contributions

AN conceived, designed, and initiated the study, performed all the studies of neurons in culture, and quantified the expression levels of complex I-V in mouse brain samples. He also led the entire study, wrote the first draft of the paper and along with GSB, served as the main editor of the original text. SRA and PR did optimization of the 2P-FLIM assay in live mouse brain. VS and PR performed 2P-FLIM experiments in the live mouse brain. HW and VS analyzed 2P-FLIM experiments in the live cells and mouse brains. AP provided extensive technical support for 2P-FLIM. EP and TK took care of the animal colony, performed q-PCR, and along with AN and VS coordinated live mouse experiments. ZW, ZF, and SH performed and analyzed experiments involving MP-PAM. GSB analyzed data and provided intellectual feedback throughout the study. All other authors read, edited, and approved the submission of the manuscript.

## Legends of Supplemental Figures

**Supplemental Figure 1. WT and APP^SAA^ mice cortex express similar levels of electron transport chain (ETC) subunits in the cortex.**

A, Cortical protein samples were separated by SDSPAGE followed by western blots of the indicated proteins. The blots shown are representative of 3 independent experiments. B-F, The expression levels of each of the indicated subunits of the electron transport chain (CI-CV, or Complex I-V) were normalized to expression of TOM40 (mitochondrial marker) detected on each sample. Bar graphs shows the quantification obtained using 3 mice/group/genetic background and analyzed using Student’s two-tailed paired t-test.

**Supplemental Figure 2. NiMA is also disrupted in female APP^SAA/+^ brain.**

A, 2P-FLIM imaging of WT female cerebral cortex through an open-skull window before and after topical application of amino acids (R+L). An increase in a2% was observed for several minutes indicating upregulation of mitochondrial activity. Statistical analyses were performed using Student’s two-tailed paired t-test. 3 females were used in these experiments. B-A similar experiment was performed using a female APP^SAA/+^ mouse cerebral cortex. Like we observed in males, NiMA is also blocked in the 4-month-old females. Graphs show the quantification of 3 independent experiments. Statistical analyses were performed using Student’s two-tailed unpaired t-test. Comparisons between time 0 and 30, 60 and 80 minutes in histograms resulted in p values >0.05.

**Supplemental Figure 3. Lithium stimulates mitochondrial activity in human neurons in culture and WT live mouse brain.**

A, Neuron cultures derived from the RenCell VM line of human neuronal progenitors were differentiated for 20 days. Then, one culture was left untreated (control) while others were treated with LiCl at the indicated concentrations for ∼80 minutes. 2P-FLIM analysis revealed that inhibition of GSK3β increased the fraction of bound NADH, indicating a rise in mitochondrial respiration. Statistical analyses were performed using Student’s two-tailed paired t-test. The experiment is representative of 3 independent assays. B, LiCl efficiency was tested by western blots of protein samples from the same experiments used for 2P-FLIM. LiCl increased phosphorylation of the inhibitory site Ser-9 on GSK3β by ∼50% without affecting the expression of the mitochondrial transcription factor PGC1α. The experiment is representative of 3 independent assays.

## Notes

### Competing Interest Statement

The authors have declared no competing interest.

## References

[1] Wang W, Zhao F, Ma X, Perry G, Zhu X. Mitochondria dysfunction in the pathogenesis of Alzheimer’s disease: Recent advances. Mol Neurodegener 2020. 10.1186/s13024-020-00376-6.

[2] Hardy JA, Higgins GA. Alzheimer’s disease: the amyloid cascade hypothesis. Science 1992;256:184–5.

[3] Herrup K. The case for rejecting the amyloid cascade hypothesis. Nat Neurosci 2015;18:794–9. 10.1038/nn.4017.

[4] Musiek ES, Holtzman DM. Three dimensions of the amyloid hypothesis: Time, space and “wingmen.” Nat Neurosci 2015. 10.1038/nn.4018.

[5] Reardon S. FDA approves Alzheimer’s drug lecanemab amid safety concerns. Nature 2023;613:227–8. 10.1038/d41586-023-00030-3.

[6] Hardy J, Selkoe DJ. The Amyloid Hypothesis of Alzheimer’s Disease: Progress and Problems on the Road to Therapeutics. Science (1979) 2002;297:353–6. 10.1126/science.1072994.

[7] Norambuena A, Wallrabe H, Cao R, Wang DB, Silva A, Svindrych Z, et al. A novel lysosome-to-mitochondria signaling pathway disrupted by amyloid-β oligomers. EMBO J 2018:e100241. 10.15252/embj.2018100241.

[8] Saxton RA, Sabatini DM. mTOR Signaling in Growth, Metabolism, and Disease. Cell 2017;168:960–76. 10.1016/j.cell.2017.02.004.

[9] Norambuena A, Wallrabe H, McMahon L, Silva A, Swanson E, Khan SSS, et al. mTOR and neuronal cell cycle reentry: How impaired brain insulin signaling promotes Alzheimer’s disease. Alzheimer’s and Dementia 2017;13:152–67. 10.1016/j.jalz.2016.08.015.

[10] Norambuena A, Sun X, Wallrabe H, Cao R, Sun N, Pardo E, et al. SOD1 mediates lysosome-to-mitochondria communication and its dysregulation by amyloid-β oligomers. Neurobiol Dis 2022;169:105737. 10.1016/j.nbd.2022.105737.

[11] Hooper C, Killick R, Lovestone S. The GSK3 hypothesis of Alzheimer’s disease. J Neurochem 2008;104:1433–9. 10.1111/j.1471-4159.2007.05194.x.

[12] Hanger DP, Hughes K, Woodgett JR, Brion J-P, Anderton BH. Glycogen synthase kinase-3 induces Alzheimer’s disease-like phosphorylation of tau: Generation of paired helical filament epitopes and neuronal localisation of the kinase. Neurosci Lett 1992;147:58–62. 10.1016/0304-3940(92)90774-2.

[13] Muyllaert D, Kremer A, Jaworski T, Borghgraef P, Devijver H, Croes S, et al. Glycogen synthase kinaseJ3β, or a link between amyloid and tau pathology? Genes Brain Behav 2008;7:57–66. 10.1111/j.1601-183X.2007.00376.x.

[14] Vossel KA, Xu JC, Fomenko V, Miyamoto T, Suberbielle E, Knox JA, et al. Tau reduction prevents Aβ-induced axonal transport deficits by blocking activation of GSK3β. Journal of Cell Biology 2015;209:419–33. 10.1083/jcb.201407065.

[15] Hedgepeth CM, Conrad LJ, Zhang J, Huang H-C, Lee VMY, Klein PS. Activation of the Wnt Signaling Pathway: A Molecular Mechanism for Lithium Action. Dev Biol 1997;185:82–91. 10.1006/dbio.1997.8552.

[16] Stambolic V, Ruel L, Woodgett JR. Lithium inhibits glycogen synthase kinase-3 activity and mimics Wingless signalling in intact cells. Current Biology 1996;6:1664–9. 10.1016/S0960-9822(02)70790-2.

[17] Martin SA, Souder DC, Miller KN, Clark JP, Sagar AK, Eliceiri KW, et al. GSK3β Regulates Brain Energy Metabolism. Cell Rep 2018;23:1922–1931.e4. 10.1016/j.celrep.2018.04.045.

[18] Xia D, Lianoglou S, Sandmann T, Calvert M, Suh JH, Thomsen E, et al. Novel App knock-in mouse model shows key features of amyloid pathology and reveals profound metabolic dysregulation of microglia. Mol Neurodegener 2022;17:41. 10.1186/s13024-022-00547-7.

[19] Choi SH, Kim YH, Hebisch M, Sliwinski C, Lee S, D’Avanzo C, et al. A three-dimensional human neural cell culture model of Alzheimer’s disease. Nature 2014;515:274–8. https://doi.org/q.

[20] Cao R, Li J, Ning B, Sun N, Wang T, Zuo Z, et al. Functional and oxygen-metabolic photoacoustic microscopy of the awake mouse brain. Neuroimage 2017;150:77–87. 10.1016/j.neuroimage.2017.01.049.

[21] Lakowicz JR. Principles of Fluorescence Spectroscopy Principles of Fluorescence Spectroscopy. 2006. 10.1007/978-0-387-46312-4.

[22] Blacker TS, Mann ZF, Gale JE, Ziegler M, Bain AJ, Szabadkai G, et al. Separating NADH and NADPH fluorescence in live cells and tissues using FLIM. Nat Commun 2014;5. 10.1038/ncomms4936.

[23] Hoyer S, Oesterreich K, Wagner O. Glucose metabolism as the site of the primary abnormality in early-onset dementia of Alzheimer type? J Neurol 1988;235:143–8. 10.1007/BF00314304.

[24] Ding S, Wu TYH, Brinker A, Peters EC, Hur W, Gray NS, et al. Synthetic small molecules that control stem cell fate. Proceedings of the National Academy of Sciences 2003;100:7632–7. 10.1073/pnas.0732087100.

[25] Sutherland C, Leighton IA, Cohen P. Inactivation of glycogen synthase kinase-3 β by phosphorylation: new kinase connections in insulin and growth-factor signalling. Biochemical Journal 1993;296:15–9. 10.1042/bj2960015.

[26] De Felice FG, Gonçalves RA, Ferreira ST. Impaired insulin signalling and allostatic load in Alzheimer disease. Nat Rev Neurosci 2022;23:215–30. 10.1038/s41583-022-00558-9.

[27] Zhao W, De Felice FG, Fernandez S, Chen H, Lambert MP, Quon MJ, et al. Amyloid beta oligomers induce impairment of neuronal insulin receptors. The FASEB Journal 2008;22:246–60. 10.1096/fj.06-7703com.

[28] Gordon BA, Blazey TM, Su Y, Hari-Raj A, Dincer A, Flores S, et al. Spatial patterns of neuroimaging biomarker change in individuals from families with autosomal dominant Alzheimer’s disease: a longitudinal study. Lancet Neurol 2018;17:241–50. 10.1016/S1474-4422(18)30028-0.

[29] Croteau E, Castellano CA, Fortier M, Bocti C, Fulop T, Paquet N, et al. A cross-sectional comparison of brain glucose and ketone metabolism in cognitively healthy older adults, mild cognitive impairment and early Alzheimer’s disease. Exp Gerontol 2018;107:18–26. 10.1016/j.exger.2017.07.004.

[30] Crane PK, Walker R, Hubbard RA, Li G, Nathan DM, Zheng H, et al. Glucose Levels and Risk of Dementia. New England Journal of Medicine 2013;369:540–8. 10.1056/NEJMoa1215740.

[31] Kapogiannis D, Mattson MP. Disrupted energy metabolism and neuronal circuit dysfunction in cognitive impairment and Alzheimer’s disease. Lancet Neurol 2011;10:187–98. 10.1016/S1474-4422(10)70277-5.

[32] Yassine HN, Self W, Kerman BE, Santoni G, Navalpur Shanmugam N, Abdullah L, et al. Nutritional metabolism and cerebral bioenergetics in Alzheimer’s disease and related dementias. Alzheimer’s & Dementia 2023;19:1041–66. 10.1002/alz.12845.

[33] Stoica L, Zhu PJ, Huang W, Zhou H, Kozma SC, Costa-Mattioli M. Selective pharmacogenetic inhibition of mammalian target of Rapamycin complex I (mTORC1) blocks long-term synaptic plasticity and memory storage. Proceedings of the National Academy of Sciences 2011;108:3791–6. 10.1073/pnas.1014715108.

[34] Aman Y, Schmauck-Medina T, Hansen M, Morimoto RI, Simon AK, Bjedov I, et al. Autophagy in healthy aging and disease. Nat Aging 2021;1:634–50. 10.1038/s43587-021-00098-4.

[35] Bartolomé A, García-Aguilar A, Asahara S-I, Kido Y, Guillén C, Pajvani UB, et al. MTORC1 Regulates both General Autophagy and Mitophagy Induction after Oxidative Phosphorylation Uncoupling. Mol Cell Biol 2017;37. 10.1128/MCB.00441-17.

[36] Fang EF, Hou Y, Palikaras K, Adriaanse BA, Kerr JS, Yang B, et al. Mitophagy inhibits amyloid-β and tau pathology and reverses cognitive deficits in models of Alzheimer’s disease. Nat Neurosci 2019;22:401–12. 10.1038/s41593-018-0332-9.

[37] Oddo S. The role of mTOR signaling in Alzheimer disease. Frontiers in Bioscience 2012;S4:310. 10.2741/s310.

[38] Caccamo A, De Pinto V, Messina A, Branca C, Oddo S. Genetic reduction of mammalian target of rapamycin ameliorates Alzheimer’s disease-like cognitive and pathological deficits by restoring hippocampal gene expression signature. J Neurosci 2014;34:7988–98. 10.1523/JNEUROSCI.0777-14.2014.

[39] Domínguez JM, Fuertes A, Orozco L, del Monte-Millán M, Delgado E, Medina M. Evidence for Irreversible Inhibition of Glycogen Synthase Kinase-3β by Tideglusib. Journal of Biological Chemistry 2012;287:893–904. 10.1074/jbc.M111.306472.

[40] del Ser T, Steinwachs KC, Gertz HJ, Andrés M V., Gómez-Carrillo B, Medina M, et al. Treatment of Alzheimer’s Disease with the GSK-3 Inhibitor Tideglusib: A Pilot Study. Journal of Alzheimer’s Disease 2012;33:205–15. 10.3233/JAD-2012-120805.

[41] Lovestone S, Boada M, Dubois B, Hüll M, Rinne JO, Huppertz H-J, et al. A Phase II Trial of Tideglusib in Alzheimer’s Disease. Journal of Alzheimer’s Disease 2015;45:75–88. 10.3233/JAD-141959.

[42] Li Y, Rinne JO, Mosconi L, Pirraglia E, Rusinek H, DeSanti S, et al. Regional analysis of FDG and PIB-PET images in normal aging, mild cognitive impairment, and Alzheimer’s disease. Eur J Nucl Med Mol Imaging 2008;35:2169–81. 10.1007/s00259-008-0833-y.

[43] Rabinovici GD, Furst AJ, Alkalay A, Racine CA, O’Neil JP, Janabi M, et al. Increased metabolic vulnerability in early-onset Alzheimer’s disease is not related to amyloid burden. Brain 2010;133:512–28. 10.1093/brain/awp326.

[44] Furst AJ, Rabinovici GD, Rostomian AH, Steed T, Alkalay A, Racine C, et al. Cognition, glucose metabolism and amyloid burden in Alzheimer’s disease. Neurobiol Aging 2012;33:215–25. 10.1016/j.neurobiolaging.2010.03.011.

[45] Cohen AD, Price JC, Weissfeld LA, James J, Rosario BL, Bi W, et al. Basal Cerebral Metabolism May Modulate the Cognitive Effects of Aβ in Mild Cognitive Impairment: An Example of Brain Reserve. The Journal of Neuroscience 2009;29:14770–8. 10.1523/JNEUROSCI.3669-09.2009.

[46] Edison P, Archer HA, Hinz R, Hammers A, Pavese N, Tai YF, et al. Amyloid, hypometabolism, and cognition in Alzheimer disease. Neurology 2007;68:501–8. 10.1212/01.wnl.0000244749.20056.d4.

[47] Lowe VJ, Weigand SD, Senjem ML, Vemuri P, Jordan L, Kantarci K, et al. Association of hypometabolism and amyloid levels in aging, normal subjects. Neurology 2014;82:1959–67. 10.1212/WNL.0000000000000467.

[48] Engler H, Forsberg A, Almkvist O, Blomquist G, Larsson E, Savitcheva I, et al. Two-year follow-up of amyloid deposition in patients with Alzheimer’s disease. Brain 2006;129:2856–66. 10.1093/brain/awl178.

[49] Altmann A, Ng B, Landau SM, Jagust WJ, Greicius MD. Regional brain hypometabolism is unrelated to regional amyloid plaque burden. Brain 2015;138:3734–46. 10.1093/brain/awv278.

[50] LEE H, CASADESUS G, ZHU X, TAKEDA A, PERRY G, SMITH MA. Challenging the Amyloid Cascade Hypothesis: Senile Plaques and AmyloidJβ as Protective Adaptations to Alzheimer Disease. Ann N Y Acad Sci 2004;1019:1–4. 10.1196/annals.1297.001.

[51] Cuajungco MP, Goldstein LE, Nunomura A, Smith MA, Lim JT, Atwood CS, et al. Evidence that the β-Amyloid Plaques of Alzheimer’s Disease Represent the Redox-silencing and Entombment of Aβ by Zinc. Journal of Biological Chemistry 2000;275:19439–42. 10.1074/jbc.C000165200.

[52] Malkov A, Popova I, Ivanov A, Jang S-S, Yoon SY, Osypov A, et al. Aβ initiates brain hypometabolism, network dysfunction and behavioral abnormalities via NOX2-induced oxidative stress in mice. Commun Biol 2021;4:1054. 10.1038/s42003-021-02551-x.

[53] Bloom GS. Amyloid-β and Tau: the Trigger and Bullet in Alzheimer’s Disease Pathogenesis. JAMA Neurol 2014;71:505. 10.1001/jamaneurol.2013.5847.

[54] Selkoe DJ. Alzheimer’s Disease Is a Synaptic Failure. Science (1979) 2002;298:789–91. 10.1126/science.1074069.

[55] Bloom GS, Norambuena A. Dysregulation of <scp>mTOR</scp> by tau in Alzheimer’s disease. Cytoskeleton 2023. 10.1002/cm.21782.

