## Supplementary figures and images for "Disrupted mitochondrial response to nutrients is a presymptomatic event in the cortex of the APP^SAA^ knock-in mouse model of Alzheimer’s disease"

### Supplemental Figure 1

#### Supplemental Figure 1
##### Norambuena et al.

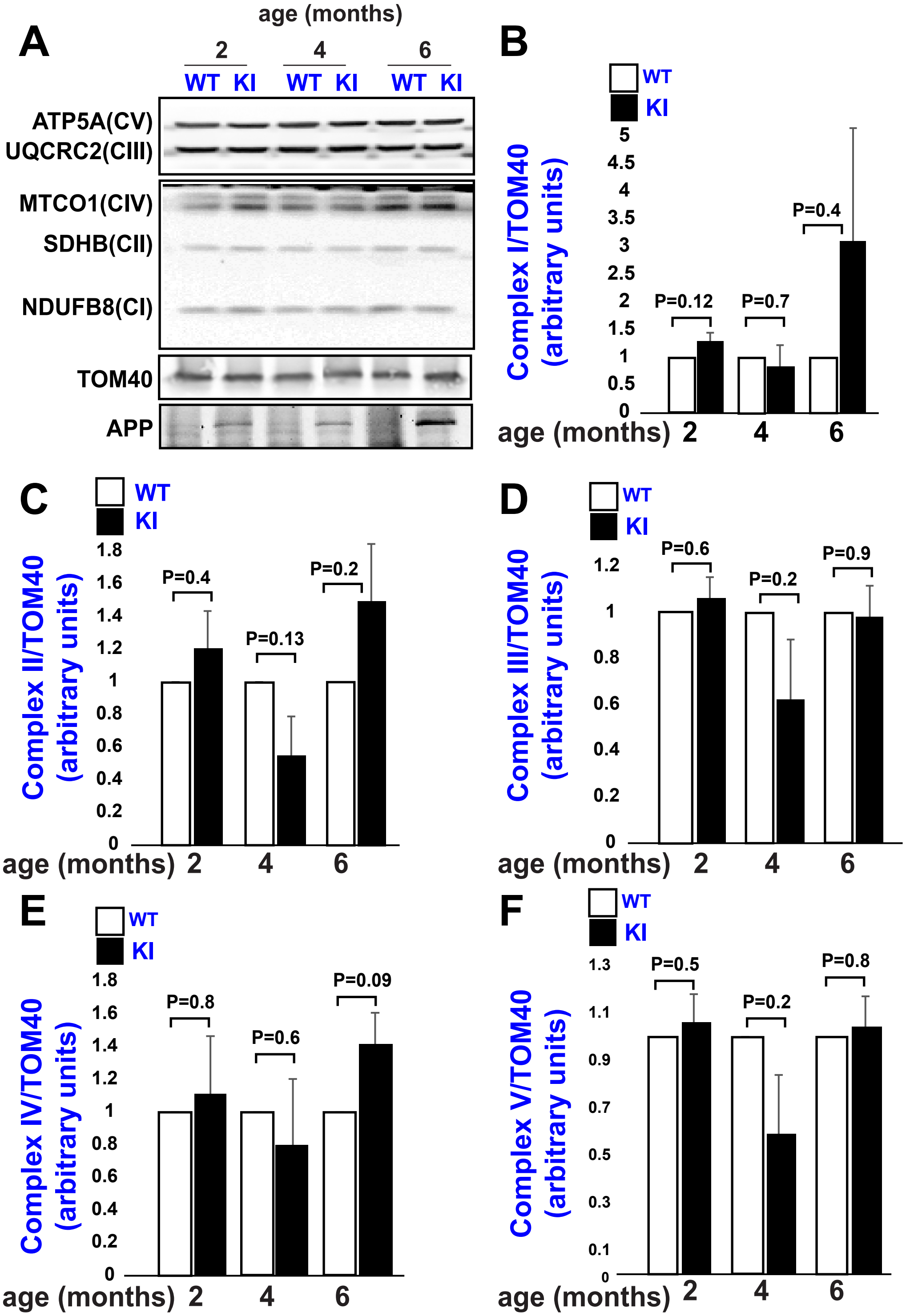

### Supplemental Figure 2

**Supplemental Figure 2**
**Norambuena et al.**

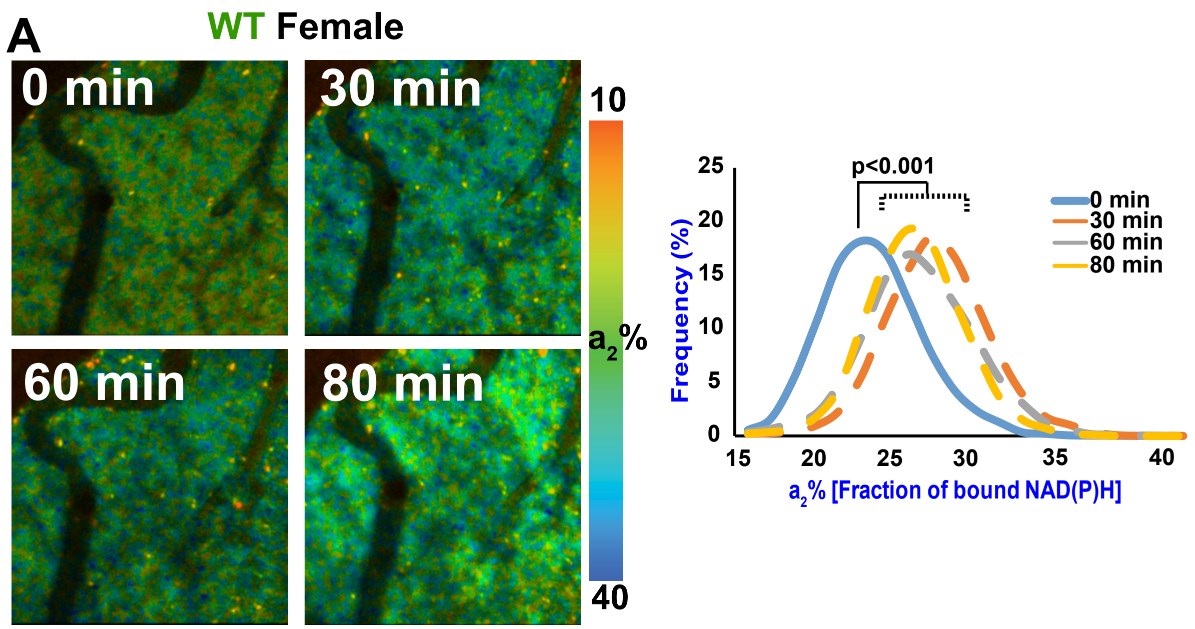


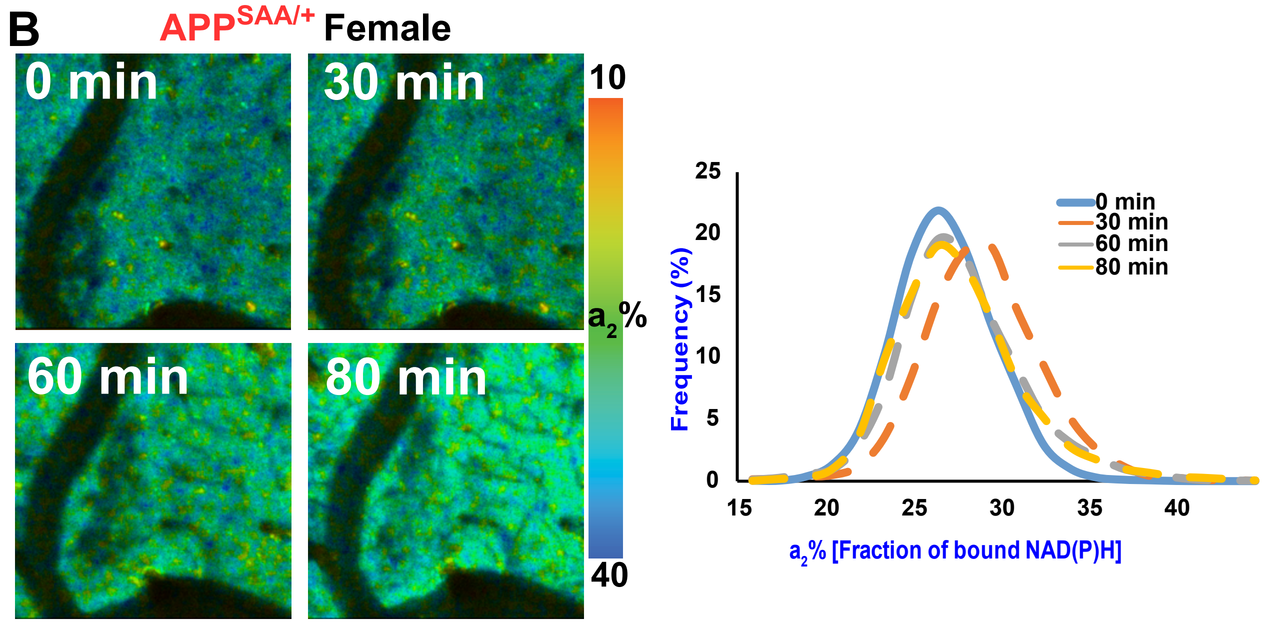

### Supplemental Figure 3

#### Supplemental Figure 3
Norambuena et al.

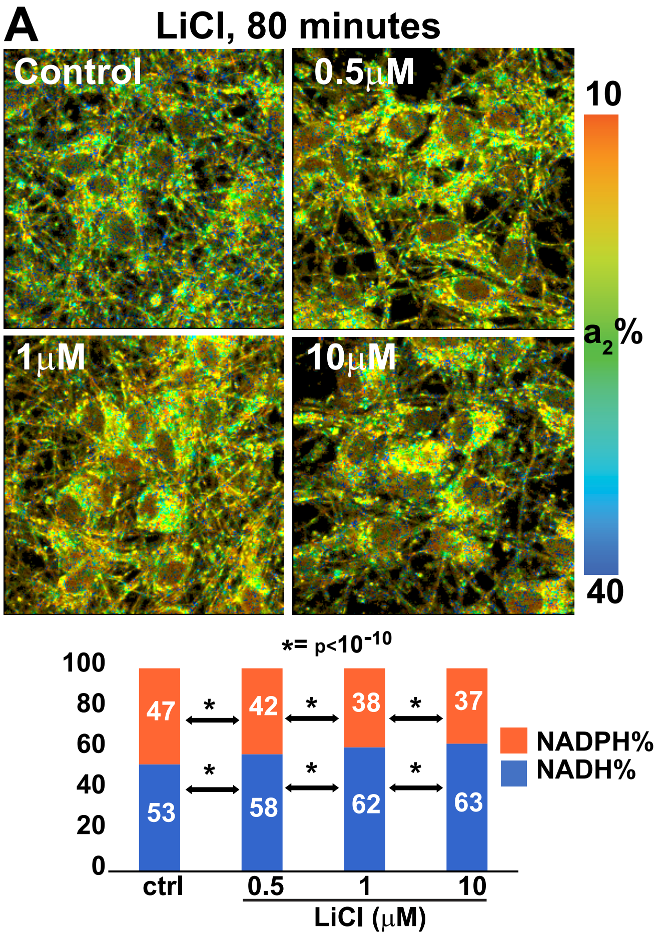


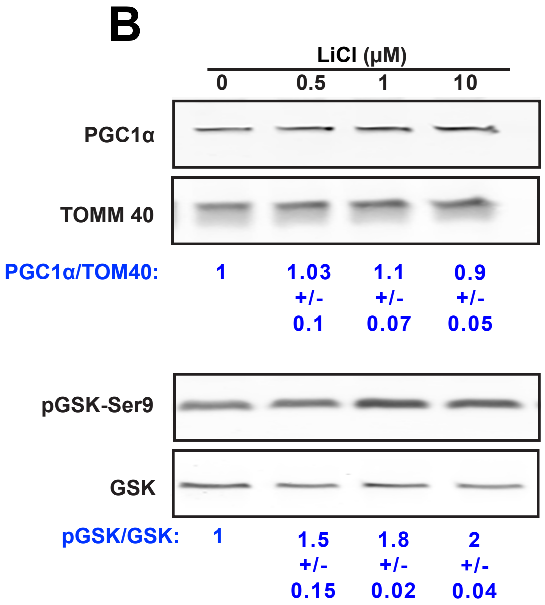
